# Free ammonia pretreatment for anaerobic sludge digestion reduces the spread of antibiotic resistance

**DOI:** 10.1101/2020.09.11.291054

**Authors:** Zehao Zhang, Huan Liu, Qilin Wang

## Abstract

Sludge from the wastewater treatment plants (WWTPs) has been recognized as a reservoir of antibiotic resistance genes (ARGs). Free ammonia (FA, i.e. NH_3_-N) pretreatment has been demonstrated to be able to enhance anaerobic digestion, which is a widely used method for sludge treatment. However, the effect of combined FA pretreatment and anaerobic digestion on the fate of ARGs is still unknown. This study demonstrated for the first time that combined FA pretreatment (420 mg NH_3_-N/L for 24 h) and anaerobic digestion could reduce the abundances of the tested ARGs by 0.06 log_10_ gene copies/g TS (total solids) compared with the anaerobic digestion alone. Specifically, the experimental results showed that combined FA pretreatment and anaerobic digestion reduced the abundances of *aac(6’)-Ib-cr, blaTEM, sul2, tetA*, *tetB* and *tetX* by 0.07, 0.37, 0.09, 0.32, 0.24 and 0.59 log_10_ gene copies/g TS compared with anaerobic digestion alone. In contrast, combined FA pretreatment and anaerobic digestion slightly increased the abundance of *tetG* by 0.05 log_10_ gene copies/g TS compared with anaerobic digestion alone. In addition, FA pretreatment did not significantly affect the abundance of *sul1 and tetM* during anaerobic digestion. This study revealed that FA pretreatment for anaerobic digestion could potentially reduce the spread of antibiotic resistance from the sludge to soil (while agriculture reuse is used as the sludge disposal method), thereby protecting the environment and human health.

## Introduction

Antibiotics have been produced and applied in medical care, poultry farming and aquaculture to promote human health and agricultural production for more than 70 years (Xue et al., 2019). However, antibiotic resistance caused by the intensive use of antibiotics poses a global threat to public well-being (Pei et al., 2016). For instance, the Center for Disease Control and Prevention reported the death of more than 35,000 people in America each year owing to antibiotic resistance.

Antibiotic resistance genes (ARGs) widely disseminate in bacterial communities through vertical gene transfer and horizontal gene transfer (Shao et al., 2018; Xue et al., 2019; Zhang et al., 2018). The VGT means antibiotic resistance bacteria (ARB) pass on ARGs through cell reproduction (Shao et al., 2018). The horizontal gene transfer means non-ARB obtain one or multiple ARGs from ARB or environments (Shao et al., 2018; Xue et al., 2019; Zhang et al., 2018).

Sludge in wastewater treatment plants (WWTPs) is recognized as a reservoir and environmental supplier for ARGs and ARB (Heuer et al., 2011; Karkman et al., 2018; Smets and Barkay, 2005; Zhu et al., 2013). The antibiotics from domestic households and hospitals enter into the WWTPs, which causes the occurrence of ARGs in various wastewater treatment bacteria (Aminov, 2011; Karkman et al., 2018). More than 99% of these ARGs will then finally accumulate in sludge (Xue et al., 2019; Yang et al., 2014). For instance, tetracyclines (*tet*) are spectrum antibiotics, which are commonly used in humans, livestock, and aquaculture (Martınez et al., 2015; Wang et al., 2016). These antibiotics have caused the occurrence of significant amounts of *tet* resistance genes in sludge. It was reported that the abundance of *tetA* and *tetQ* in sludge could reach 10^8^-10^9^ copies/g-TS (total solids) and 10^4^-10^7^ copies/g-TS, respectively (Auerbach et al., 2007). Agricultural reuse is a common sludge disposal method. For example, more than 67% of sludge is reused in agriculture in Australia (Australian Water Association, 2020). This may lead to the spread of ARGs from sludge to the local environment, thereby increasing the risk of sludge reuse to human health.

Anaerobic digestion is a typical sludge treatment method because it achieves sludge reduction (Batstone et al., 2002). Recently, it is reported that anaerobic digestion can also affect the fate of some types of ARGs. For instance, Pei et al. (2016) found that anaerobic digestion could decrease the total abundance of *tetA*, *tetG*, *tetQ*, *tetQ* and *tetX* by up to 1.00 log_10_ gene copies/g TS. Anaerobic digestion is often limited by the poor biodegradability of the sludge. Therefore, several pretreatment methods have been developed to enhance sludge biodegradability, thereby enhancing sludge reduction. For instance, it has been reported that free ammonia (FA, i.e., NH_3_) pretreatment at 420 mg N/L for 24 h enhanced sludge biodegradability by 20% (Wei et al., 2017). The FA pretreatment technology relies on a renewable material (i.e. FA) from wastewater, making this technology quite promising.

Recently, it has been found that some pretreatment technologies affected the fate of ARGs. For instance, Pei et al. (2016) showed that the thermal hydrolysis pretreatment could enhance the reduction of five *tet* resistance genes by 0.5 log copies/g-TS in anaerobic digestion. However, it is still unknown how the FA pretreatment technology will affect the fate of ARGs in anaerobic digestion.

This study aimed to assess the effect of FA pretreatment on the spread of antibiotic resistance during anaerobic sludge digestion. Nine ARGs were selected to quantify in this study, which were *aac(6’)-Ib-cr*, *blaTEM, sul1, sul2, tetA, tetB, tetG, tetM* and *tetX*.

## Materials and methods

### Secondary sludge and inoculum

The secondary sludge was collected from the thickener of a wastewater treatment plant (WWTP) conducting biological nitrogen and phosphorus removal. The WWTP has a sludge retention time (SRT) of 12 - 16 d. The inoculum was collected from a mesophilic anaerobic digester treating a mixture of primary and secondary sludges in the WWTP from which secondary sludge was collected. The mesophilic anaerobic digester has an SRT of 15 - 18 d.

### Pretreatment of secondary sludge with FA

Batch experiments were conducted to evaluate the impact of FA pretreatment on the abundances of detected ARGs. 1 L of secondary sludge was evenly distributed into two batch reactors. For FA pretreatment, a certain amount of ammonium stock solution (3.0 M) was added to one of the two batch reactors to obtain a total ammonia nitrogen (NH_4_^+^-N + NH_3_-N) concentration of 500 mg N/L. pH was adjusted and maintained at 10.0 ± 0.1 using NaOH solution. The pretreatment was conducted in a temperature-controlled room (25 ℃) and lasted for 24 h. The total ammonia nitrogen, pH and the temperature collectively resulted in an FA concentration of 420 mg NH_3_-N/L, which was determined by the formula S_(NH4−N+NH_3_-N)_ × 10^pH^/(K_b_/K_w_+10^pH^). The S_(NH4−N+NH_3_-N)_ is the total ammonia nitrogen concentration. The K_b_/K_w_ is equal to e^6,344/(273+T)^ (Anthonisen et al., 1976). This ammonia concentration (i.e. 420 mg NH_3_-N/L) was selected based on our previous tests (Wei et al., 2017), which demonstrated that FA pretreatment at 420 mg NH_3_-N/L for 24 h led to the highest methane production with a large economic advantage. The other batch reactor was also set up without ammonium addition or pH control. This reactor served as a control. The sludge samples were taken both before and after pretreatment for the determination ARGs using real-time quantitative PCR (RT-qPCR) to be described below.

### Anaerobic digestion tests

Anaerobic digestion tests were performed to evaluate the effect of combined FA pretreatment with anaerobic digestion on the abundances of detected ARGs. The serum vials (160 mL) with a working volume of 100 mL were used to carry out the anaerobic digestion tests. Both the inoculum and the secondary sludge were added into every serum vial, resulting in a VS based inoculum to sludge ratio of approximately 2.0. The vials were flushed with helium gas for 2 min (1 L/min) to ensure an anaerobic condition. After that, a rubber stopper with an aluminum crimp cap was used to seal the vials, which were then put in an incubator operated at 37 ℃. Triplicate tests were conducted. The anaerobic digestion tests lasted for 45 days. The digested sludges with and without FA pretreatment were sampled for the analysis of ARGs by *RT-qPCR*.

### DNA extraction and RT-qPCR

0.25 g of sludge of each sample was used for DNA extraction using the Fast DNA Spin Kit for Soil (MP Biomedicals, USA). 1% agarose gel electrophoresis and NanoDrop ND-1000 (NanoDrop, USA) were used to determine the quality and concentration of the extracted DNA. Triplicate extracts were conducted to get a representative DNA sample.

RT-qPCR was used for the quantification of ARGs. One aminoglycoside and fluoroquinolone resistance gene (i.e. *aac(6’)-Ib-cr*), one beta-lactamase resistance gene (i.e. *blaTEM*), two sulfonamide resistance genes (i.e. *sul1* and *sul2*) and five *tet* resistance genes (i.e. *tetA, tetB, tetG, tetM* and *tetX)* were selected in this study. PCR was conducted based on the method in Zhang et al. (2017). The primers and standard PCR conditions employed are described in the Supporting Information (Table S1). The abundances of ARGs were expressed as the gene copies divided by the TS of the sludge (i.e., gene copies per gram TS).

## Results

### Effects of FA pretreatment on the fate of ARGs

The effect of FA pretreatment on the fate of ARGs in sludge is shown in Fig. 1 and Table 1, which show FA pretreatment significantly affected the abundance of ARGs. FA pretreatment at 420 mg N/L for 24 h decreased the abundances of *blaTEM*, *sul1*, *sul2*, *tetA*, *tetB* and *tetX* by 0.10, 0.04, 0.58, 0.20 and 0.61 log_10_ copies/g-TS, respectively, which were from 7.76×10^6^, 1.64×10^9^, 6.35×10^8^, 3.40×10^7^, 1.72×10^8^ and 7.82×10^7^ gene copies/g-TS to 6.17×10^6^, 1.51×10^9^, 5.31×10^8^, 9.11×10^6^, 1.07×10^8^ and 1.93×10^7^ gene copies/g-TS, respectively. In contrast, FA pretreatment increased the abundance of *tetM* by 0.71 log_10_ copies/g-TS, which were from 4.09×10^6^ gene copies/g-TS to 2.10×10^7^ gene copies/g-TS. In terms of *aac(6’)-Ib-cr* and *tetG*, FA pretreatment did not significantly affect their abundances, which were maintained at around 3.30×10^9^ and 9.00×10^7^ gene copies/g-TS both before and after FA pretreatment. Overall, FA pretreatment at 420 mg N/L for 24 h decreased the abundance of the tested ARGs by 0.04 log_10_ copies/g-TS, which decreased from 6.01×10^9^ gene copies/g-TS to 5.47×10^9^ gene copies/g-TS. This demonstrated that FA pretreatment is effective in removing the tested ARGs.

**Table 1.** Effects of FA pretreatment on the fate of ARGs

|  | Abundances of the ARGs (log<sub>10</sub> gene copies/g-TS) |  |  |  |  |  |  |  |  | **ARGs** |
| --- | --- | --- | --- | --- | --- | --- | --- | --- | --- | --- |
|  | *aac(6')-ib-cr* | *blaTEM* | *sul1* | *sul2* | *tetA* | *tetB* | *tetG* | *tetM* | *tetX* |  |
| Untreated sludge | 9.52 | 6.89 | 9.22 | 8.8 | 7.53 | 8.23 | 7.94 | 6.61 | 7.9 | 9.78 |
| FA-pretreated sludge | 9.5 | 6.79 | 9.18 | 8.72 | 6.95 | 8.03 | 7.98 | 7.32 | 7.29 | 9.74 |
| Δ Abundance | 0.02 | 0.1 | 0.04 | 0.08 | 0.58 | 0.2 | -0.04 | -0.71 | 0.61 | 0.04 |

Note: The negative value means the target gene is increased.

**Figure 1.**
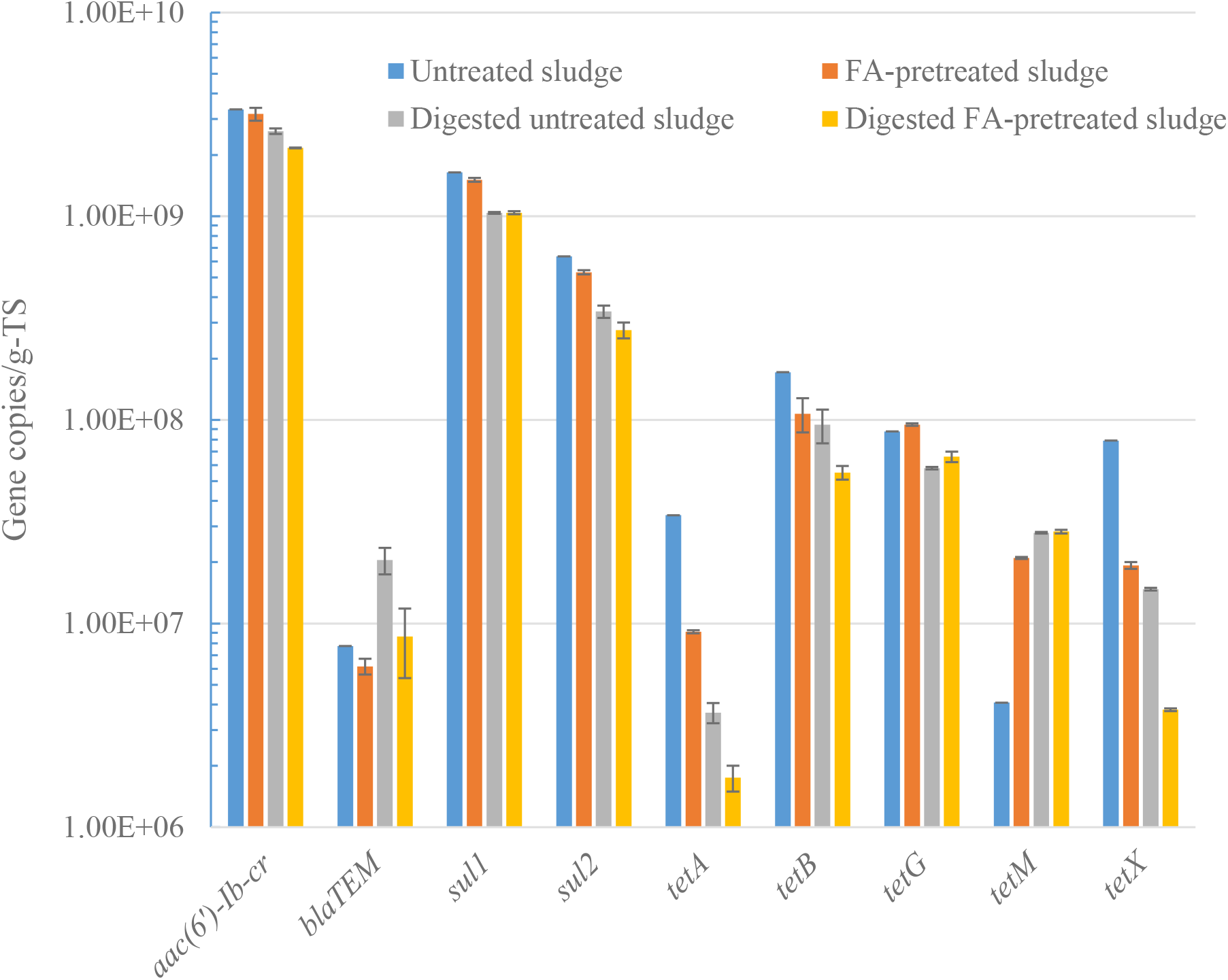
Abundances of ARGs in different sludges

### Effects of combined FA pretreatment and anaerobic digestion on the fate of ARGs

The effect of combined FA pretreatment and anaerobic digestion on the fate of ARGs is shown in Fig. 1 and Table 2. Compared with the anaerobic digestion alone, combined FA pretreatment and anaerobic digestion could further reduce the abundances of *aac(6’)-ib-cr*, *blaTEM*, *sul2*, *tetA, tetB* and *tetX* by 0.07, 0.37, 0.09, 0.32, 0.24 and 0.59 log_10_ copies/g-TS, respectively, which were from 2.62×10^9^, 2.05×10^7^, 3.40×10^8^, 3.65×10^6^, 9.45×10^7^ and 1.48×10^7^ gene copies/g-TS in the digested untreated sludge to 2.16×10^9^, 8.62×10^6^, 2.76×10^8^ 1.90×10^6^, 3.94×10^7^ and 1.10×10^7^ gene copies/g-TS in the digested FA-pretreated sludge, respectively. On the contrary, combined FA pretreatment and anaerobic digestion slightly increased the abundance of *tetG* by 0.06 log_10_ copies/g-TS compared with the anaerobic digestion alone, which was from 5.79×10^7^ gene copies/g-TS in the digested untreated sludge to 6.59×10^7^ gene copies/g-TS in the digested FA-pretreated sludge. In addition, combined FA pretreatment and anaerobic digestion did not affect the abundances of *sul1* and *tetM*, which were maintained at approximately 1.04×10^9^ and 2.80×10^7^ gene copies/g-TS in both digested untreated sludge and digested FA-pretreated sludge. In total, combined FA pretreatment and anaerobic digestion reduced the abundance of the tested ARGs by 0.06 log_10_ copies/g-TS compared with the anaerobic digestion alone, which was from 4.21×10^9^ gene copies/g-TS in the digested untreated sludge to 3.64×10^9^ gene copies/g-TS in the digested FA-pretreated sludge. Therefore, combined FA pretreatment and anaerobic digestion is effective in reducing the abundance of the tested ARGs.

**Table 2.** Effects of combined FA pretreatment and anaerobic digestion on the fate of ARGs

|  | Abundances of the ARGs (log<sub>10</sub> gene copies/g-TS) |  |  |  |  |  |  |  |  | **ARGs** |
| --- | --- | --- | --- | --- | --- | --- | --- | --- | --- | --- |
|  | *aac(6')-ib-cr* | *blaTEM* | *sul1* | *sul2* | *tetA* | *tetB* | *tetG* | *tetM* | *tetX* |  |
| Digested untreated sludge | 9.41 | 7.31 | 9.02 | 8.53 | 6.56 | 7.98 | 7.76 | 7.45 | 7.17 | 9.62 |
| Digested FA-pretreated sludge | 9.34 | 6.94 | 9.02 | 8.44 | 6.24 | 7.74 | 7.81 | 7.45 | 6.58 | 9.56 |
| Δ Abundance | 0.07 | 0.37 | 0.00 | 0.09 | 0.32 | 0.24 | -0.05 | 0.00 | 0.59 | 0.06 |

Note: The negative value means the target gene is increased.

## Conclusion

Compared with anaerobic digestion alone, combined FA pretreatment and anaerobic digestion could reduce the abundances of antibiotic resistance genes. It reduced the abundances of *aac(6’)-ib-cr*, *blaTEM*, *sul2*, *tetA, tetB* and *tetX* by 0.07, 0.37, 0.09, 0.32, 0.24 and 0.59 log_10_ copies/g-TS compared with the anaerobic digestion alone, but slightly increased the abundance of *tetG* by 0.06 log_10_ copies/g-TS with the anaerobic digestion alone. Combined FA pretreatment and anaerobic digestion did not significantly affect the abundances of *sul1* and *tetM* compared with the anaerobic digestion alone. FA pretreatment can be used as a potential method to reduce the spread of antibiotic resistance from sludge to the environment.

