## Supplemental Table 1 for "Free ammonia pretreatment for anaerobic sludge digestion reduces the spread of antibiotic resistance"

**Table S1. Primers used in this study**

| ARGs | Primer Sequences Forward | Primer Sequences Reverse | Amplicon size (bp) | Annealing temperature (℃) | References |
|---|---|---|---|---|---|
| *aac(6')-ib-cr* | TTGCGATGCTCTATGAGTGGCTA | CTCGAATGCCTGGCGTGTTT | 482 | 66 | (Jang et al., 2017) |
| *blaTEM* | ATCAGCAATAAACCAGC | CCCCGAAGAACGTTTTC | 516 | 60 | (Zhang et al., 2017) |
| *sul1* | CGCACCGGAAACATCGCTGCAC | TGAAGTTCCGCCGCAAGGCTCG | 163 | 65 | (Pei et al., 2006) |
| *sul2* | TCCGGTGGAGGCCGGTATCTGG | CGGGAATGCCATCTGCCTTGAG | 191 | 57.5 | (Pei et al., 2006) |
| *tetA* | GCTACATCCTGCTTGCCTTC | CATAGATCGCCGTGAAGAGG | 210 | 60.9 | (Jang et al., 2017) |
| *tetB* | GGTTGAGACGCAATCGAATT | AGGCTTGGAATACTGAGTGTAA | 206 | 52.9 | (Jang et al., 2017) |
| *tetG* | GCAGAGCAGGTCGCTGG | CCYGCAAGAGAAGCCAGAAG | 134 | 64.2 | (Zhang et al., 2017) |
| *tetM* | CCYGCAAGAGAAGCCAGAAG | ACAGAAAGCTTATTATATAAC | 171 | 55 | (Zhang et al., 2017) |
| *tetX* | CAATAATTGGTGGTGGACCC | TTCTTACCTTGGACATCCCG | 468 | 60 | (Zhang et al., 2017) |
